# The regulatory functions of ESX-1 substrates, EspE and EspF, are separable from secretion

**DOI:** 10.1101/2024.07.05.602283

**Authors:** Rebecca J. Prest, Konstantin V. Korotkov, Patricia A. Champion

## Abstract

Pathogenic mycobacteria are a significant global health burden. The ESX-1 secretion system is essential for mycobacterial pathogenesis. The secretion of ESX-1 substrates is required for phagosomal lysis, which allows the bacteria to enter the macrophage cytoplasm, induce a Type I IFN response, and spread to new host cells. EspE and EspF are dual-functioning ESX-1 substrates. Inside the mycobacterial cell, they regulate transcription of ESX- 1-associated genes. Following secretion, EspE and EspF are essential for lytic activity. The link between EspE/F secretion and regulatory function has not been investigated. We investigated the relationship between EspE and EspF using molecular genetics in *Mycobacterium marinum,* a non-tuberculous mycobacterial species that serves as an established model for ESX-1 secretion and function in *M. tuberculosis.* Our data support that EspE and EspF, which require each other for secretion, directly interact. Disruption of the predicted protein-protein interaction abrogates hemolytic activity and secretion but does not impact their gene regulatory activities in the mycobacterial cell. In addition, we predict a direct protein-protein interaction between the EsxA/EsxB heterodimer and EspF. Our data support that the EspF/EsxA interaction is also required for hemolytic activity and EspE secretion. Our study sheds light on the intricate molecular mechanisms governing the interactions between ESX-1 substrates, regulatory function and ESX-1 secretion, moving the field forward.

**Importance:** Tuberculosis (TB), caused by *Mycobacterium tuberculosis,* is a historical and pervasive disease responsible for millions of deaths annually. The rise of antibiotic and treatment-resistant TB, as well as the rise of infection by non-tuberculous mycobacterial species, calls for better understanding of pathogenic mycobacteria. The ESX-1 secreted substrates EspE and EspF are required for mycobacterial virulence, and may be responsible for phagosomal lysis. This study focuses on the mechanism of EspE and EspF secretion from the mycobacterial cell.

## Introduction

*Mycobacterium tuberculosis* is a global health burden that continues to infect and kill millions of people annually (1, 2). *M. tuberculosis* replicates and spreads cell-to-cell through host tissue in a cycle of uptake into host phagocytes, lysis of the phagosome, and subsequent host cell death (3–5). Rupture of the phagosomal membrane allows mycobacterial access to the macrophage cytosol, which is crucial for mycobacterial survival in the host (6–8). The ESX-1 (ESAT-6 System 1) secretion system transports proteins to the mycobacterial surface and into the host cell that are required for lysis of the phagosomal membrane (7, 9). ESX-1 is conserved across pathogenic mycobacterial species, including the opportunistic, nontuberculous species *Mycobacterium marinum* (10). *M. marinum* causes a tuberculosis-like infection in poikilotherms, and serves as a well-established model for studying mycobacterial virulence (11).

The mechanisms leading to ESX-1 assembly in the mycobacterial cell envelope and protein substrate secretion are mainly unknown. At least 12 ESX-1 substrates are secreted from *M. marinum*, including Esx WXG-100 proteins, PE/PPE proteins, and Esp proteins (ESX-1 associated proteins) (12). We previously proposed that the ESX-1 substrates are hierarchically secreted; early substrates have a role in transporting later substrates (13). Early, or ‘group I’ substrates include PPE68, MMAR_2894, EsxA (ESAT-6), and EsxB (CFP-10) (13). EsxA and EsxB form a 1:1 heterodimer, and this interaction is required for their secretion (14–16). The C-terminus of EsxB interacts with EccCb_1_, targeting the EsxA/EsxB heterodimer for transport (17). Because EsxA and EsxB are required for secretion of the successive substrate groups (13, 18), these early substrates may directly interact with the later substrates, and facilitate their interaction with EccCb_1_ for transport out of the bacterial cytosol. Following the secretion of group I substrates, the group II substrates (EspB, EspJ and EspK) and group III substrates (EspE and EspF) are secreted (13).

The group III substrates require the secretion of both group I and group II substrates for secretion (13). *espE* and *espF* are dispensable for secretion of all other substrates (13). Yet, EspE and EspF are required for mycobacterial virulence and hemolytic activity in *M. marinum* (9). EspE and EspF require each other for secretion, as the deletion of either protein results in a loss of secretion of the other, as well as a loss of virulence and hemolytic activity (9, 13). EspE and EspF also have regulatory activity inside the mycobacterial cell (9). WhiB6 positively regulates transcription of several ESX-1 substrate genes (19–21). In the absence of either EspE or EspF, WhiB6 levels are increased, which in turn causes an accumulation of ESX-1 substrates, including EsxA and EsxB (9). We previously showed that EspE and EspF negatively regulate transcription from the *whiB6* promoter. Our data suggest that the regulatory functions of EspE and EspF are not equivalent, and that EspE and EspF function as independent regulators. We showed that constitutive expression of the *espF* gene in the Δ*espE* strain or vice versa did not restore *whiB6* transcript to WT levels in the presence of the ESX-1 membrane complex (9).

Based on the interactions of paralogous proteins (22), and our prior work, it has been proposed that EspE and EspF directly interact with each other (9). However, the interaction between EspE and EspF has not been formally tested. It is also unclear if these proposed interactions are required for secretion and regulatory function of EspE and EspF. In this study, we conduct a structure-function analysis of the EspE and EspF substrates. We generated a computational model of how EspE and EspF may interact with each other, as well as with the group I substrates EsxA and EsxB, which allowed us to predict key interaction points at the protein-protein interfaces. We created mutations that disrupt the proposed interactions between EspE, EspF, and the EsxB/EsxA substrate pair. We tested the resulting protein variants to determine the role of each interaction in regulation, secretion, and ESX-1 function.

## Results

### Computational modeling predicted protein-protein interaction of group I and group III substrates

Our prior work demonstrated that the group I substrates, EsxA and EsxB, were required for the secretion of EspE and EspF (13). We also previously showed that EsxA directly interacts with EspF (23). Therefore, we hypothesized that the EsxA/EsxB dimer would directly interact with EspE and EspF. To test this hypothesis, we used AlphaFold2 (24) to create a computational model to determine if EspE and EspF interact with one another and with the EsxA/EsxB dimer. The majority of the model, and importantly, the area of the protein-protein interfaces, were high confidence (pLDDT>90).

EsxA and EsxB are WXG100 proteins, with approximately 100 amino acids with a helix, turn, helix structural motif (25). The structure of the *M. tuberculosis* ESAT-6/CFP-10 heterodimer (EsxA/EsxB orthologs), determined by NMR (14) and X-ray crystallography (26), is reflected in our model (Fig. 1). The EsxA (yellow) and EsxB (pink) termini face opposite directions to form a four-helical bundle with solvent-exposed, flexible ends. Neither EspE nor EspF has been structurally characterized using experimental methods. AlphaFold2 structure prediction reveals that, like many ESX proteins, EspF and EspE are PE/PPE-like (27). The EspF monomer is predicted to have a PE-like structure, with a flexible N-terminus. Overall, the predicted structure of EspF (purple) shows a marked similarity to those of EsxA and EsxB (28, 29). However, EspF lacks the WXG motif (25) and the C-terminal motif that interacts with the EccCb_1_ ATPase (17, 30), arguing that this substrate is secreted via a different mechanism from the EsxA/B pair. Because the C-terminal half of EspE is predicted to be largely disordered or unstructured by both AlphaFold(31) and RoseTTAFold (32)(Fig. 1A), only the PPE-like, N-terminal, amino acids 1-198 were included in the protein-protein interaction model, omitting amino acids 199-418. The N-terminal half of the EspE monomer is predicted to be made up of four α-helices (α-I through α-IV), where EspE_α-I_ and EspE_α-II_ are short and separated by a small unstructured region, and run parallel to EspE_α-III_ and EspE_α-IV_ to form a pseudo-tri-helical bundle. The gap between EspE_α-I_ and EspE_α-II_ promotes flexibility in the helical bundle. In the protein-protein interaction model, the ‘turn’ of EspF fills in this gap, opening the interior residues of EspE_α-III_ and EspE_α-IV_ to side-chain interactions. Based on this model, we hypothesize that the interaction of EspE with EspF is largely driven by the hydrophobic effect, as both proteins contain many small hydrophobic residues.

**Figure 1.**
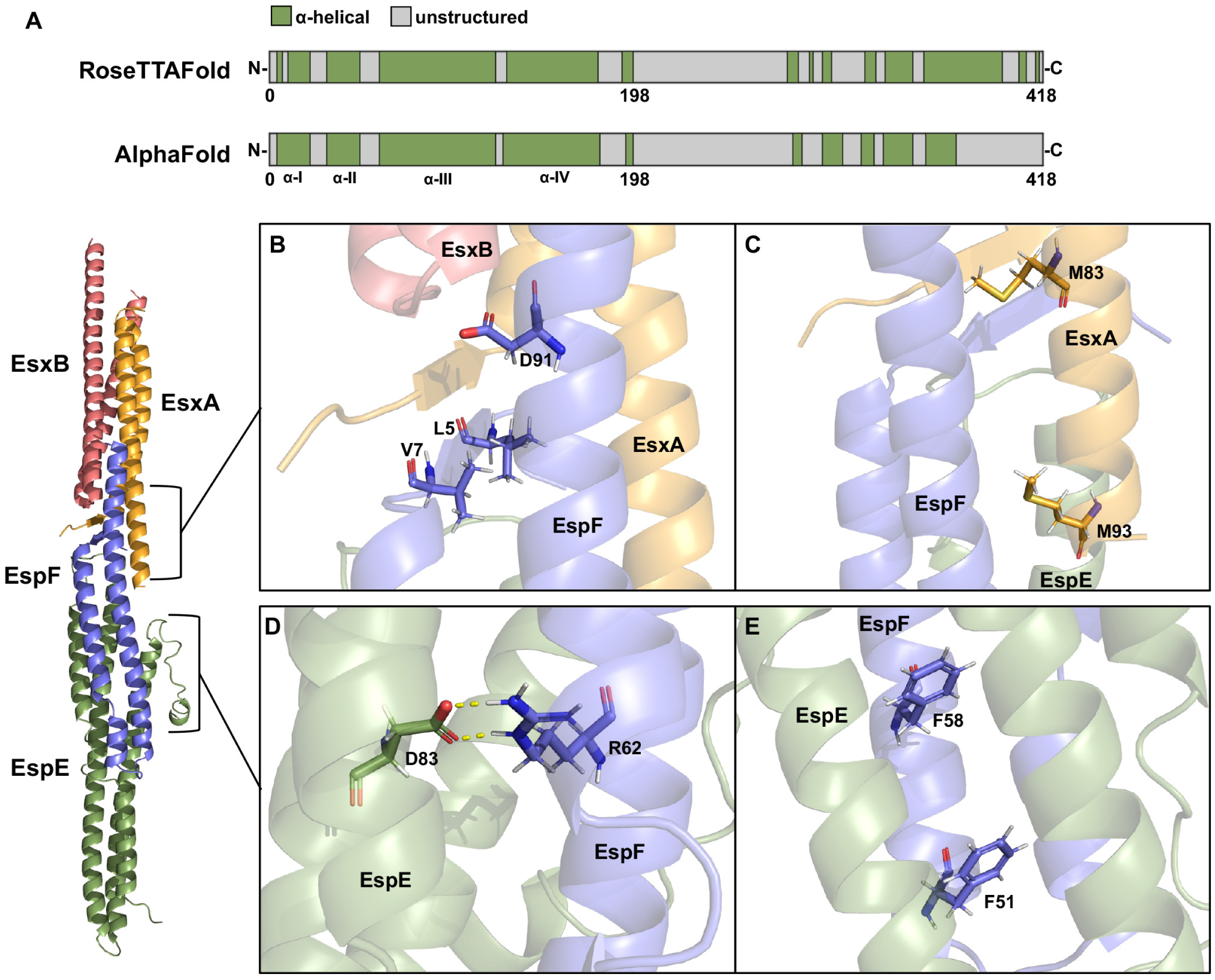
Predicted model of protein-protein interaction between group III substrates EspE and EspF with group I substrates EsxA and EsxB. A) The C-terminal half of EspE is predicted to be largely disordered or unstructured. Structural predictions of EspE (MMAR_5439) from Robetta (RoseTTAFold) (44) (top) and AlphaFold (24, 31) (bottom). B-E) Protein-protein interaction model was created using AlphaFold2 (24) and analyzed in PyMOL (The PyMOL Molecular Graphics System, Version 2.5.0 Schrödinger, LLC.). EsxB is shown in pink, EsxA in yellow, EspF in purple, and amino acids 1-198 of EspE in green. B, C) The major residues involved in the protein-protein interaction interface between EspF and EsxA. D, E) The major protein-protein interaction interface between EspE and EspF. Electrostatic interactions are represented by dashed yellow lines.

The interaction between EsxA/EsxB and EspE/EspF is largely mediated by interaction between EspF and EsxA. The flexible N-termini of EspF and EsxA are predicted to form short, 6-residue β-strands, resulting in a two-stranded, parallel, β-sheet between the two proteins (Fig. 1B). There is additional predicted side chain interaction between the C-terminal α-helices of EspF and EsxA. This leaves the flexible C-terminus of EsxB free to interact with the EccCb_1_ ATPase (17, 30). Overall, our model suggests that the secretion of EspE is mediated by the interaction of EspF and EsxA.

### EspE-EspF interaction is required for secretion and hemolytic activity

We sought to test the importance of EspE-EspF interaction in ESX-1 function. In the generated model, there is α- helix-mediated protein-protein interaction between EspE and EspF, which results in a helical bundle. We predict that the helical bundle is held together through two key regions: a salt bridge (Fig. 1 D) and a hydrophobic core (Fig. 1 E). To test the viability of the model, we generated point mutations in the *espE* and *espF* genes in the integrating plasmid p*espMEFG* containing the endogenous *espM, espE, espF,* and *espG* genes (*MMAR_5438* - *MMAR_5441*) using site-

directed mutagenesis (9). We verified each mutation using targeted Sanger sequencing and introduced the resulting plasmids into *M. marinum* strains containing unmarked deletions of the *espE* or *espF* genes (9) (Fig. 2A, B). Further, we performed RT-qPCR to verify that the mutant strains were able to produce *espE* transcript (Fig. 2E). *espE* transcript was produced in the WT strain and absent from the Δ*espE* strain. The introduction of the p*MEFG* wild-type plasmid and targeted mutations restored *espE* transcript to similar levels.

**Figure 2.**
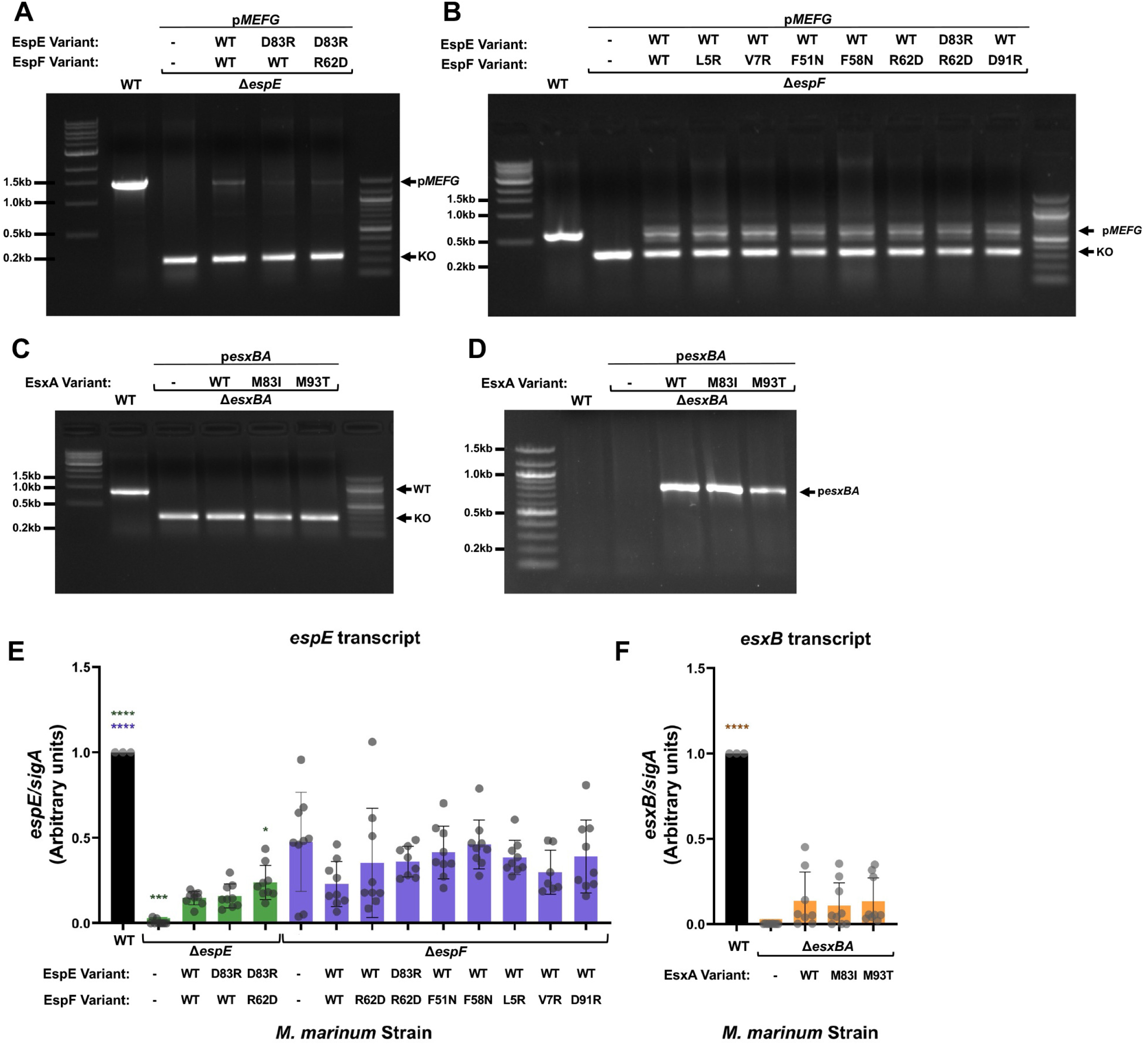
Construction of *M. marinum* strains. A) *M. marinum* strains were checked for the deletion of *espE* and the presence of the integrating plasmid p*MEFG* by PCR. WT = 1462bp, Knockout (KO) = 205bp. B) *M. marinum* strains were checked for the deletion of espF and the presence of the integrating plasmid p*MEFG* by PCR. WT = 600bp, KO = 288bp. C) *M. marinum* strains were checked for the deletion of the *esxBA* genetic locus by PCR. WT = 890bp, KO = 263bp. D) *M. marinum* strains were checked for the presence of p*esxBA* by PCR, 780bp expected. E, F) qRT-PCR analysis of *espE* (E) and *esxB* (F) transcript relative to *sigA* transcript in the indicated *M. marinum* strains. ΔΔC_T_ compared to the WT strain is shown. Data shown are three biological replicates each with three technical replicates. Statistical significance was determined using a one-way ordinary ANOVA (*P*<.0001) followed by Dunnett’s multiple comparison test relative to the Δ*espE/*p*MEFG* strain (green), the Δ*espF/*p*MEFG* strain (purple), or the Δ*esxBA/*p*esxBA* strain (yellow). * *P* = 0.0142, *** *P* = 0.0002, **** *P* < 0.0001. Error bars indicate standard deviations.

The predicted salt bridge occurs between residues EspED83 and EspFR62 (Fig. 1D). To disrupt the electrostatic attraction, we created the protein variants EspED83R and EspFR62D, which result in opposing positive or negative charges, respectively. Introduction of both protein variants in a single strain, either Δ*espE/pespED83RespFR62D* (only *espE* and *espF* alleles listed in parental plasmid *pespMEFG*) or Δ*espF/pespED83RespFR62D*, resulted in a reversal of the charges, which we hypothesized would still allow formation of the salt bridge.

Beyond the residues involved in the salt bridge, the helical bundle is largely mediated by hydrophobic side chain interactions (Fig. 1E). EspF contains several bulky hydrophobic residues, including F51 and F58, whereas EspE contains multiple small hydrophobic leucine and valine residues in this region. To disrupt the hydrophobic effect, and the predicted interaction between EspE and EspF, we replaced the phenylalanine residues of EspF with bulky hydrophilic asparagine residues to generate the EspFF51N and EspFF58N variants.

Based on the model in Figure 1, we hypothesized that disruption of areas of predicted interaction between EspE and EspF would prevent their secretion, and therefore abrogate function of ESX-1 in the mutant strains. Likewise, we predicted that the double salt bridge mutants (expressing both *espED83R* and *espFR62D*) would retain ESX-1 function. We characterized the mutations by generating cell-associated and secreted protein fractions from *M. marinum* grown in laboratory culture. Our data are summarized in Table 1. We separated the proteins using gel electrophoresis and monitored EspE levels using immunoblot analysis (Fig. 3A, S2). The β subunit of RNAP, RpoB, was used as a loading control for the cell-associated fractions, and as a lysis control for the secreted fractions. MPT-32 is a protein secreted independently of ESX-1, and serves as a loading control for both the cell-associated and secreted fractions. We are unable to detect the EspF protein by immunoblot analysis. The EspE protein was produced in and secreted from the wild-type (WT) strain as shown in Figure 3A, lane 1. EspE protein was absent from the Δ*espE* strain (lane 2). Production and secretion of EspE were restored in the complementation strain (lane 3). The EspED83R variant was detected in the presence of the WT EspF protein (lane 4) and the EspFR62D protein (lane 5), but in lower abundance than either the WT or complement strains (compare to lanes 1 and 3). The EspED83R protein was only secreted from the strain expressing the EspFR62D variant (lane 5, lower), but not from the strain expressing the WT EspF protein (lane 4, lower). From these data we conclude that disruption of the predicted salt bridge resulted in decreased production of EspE protein and loss of secretion.

**Figure 3.**
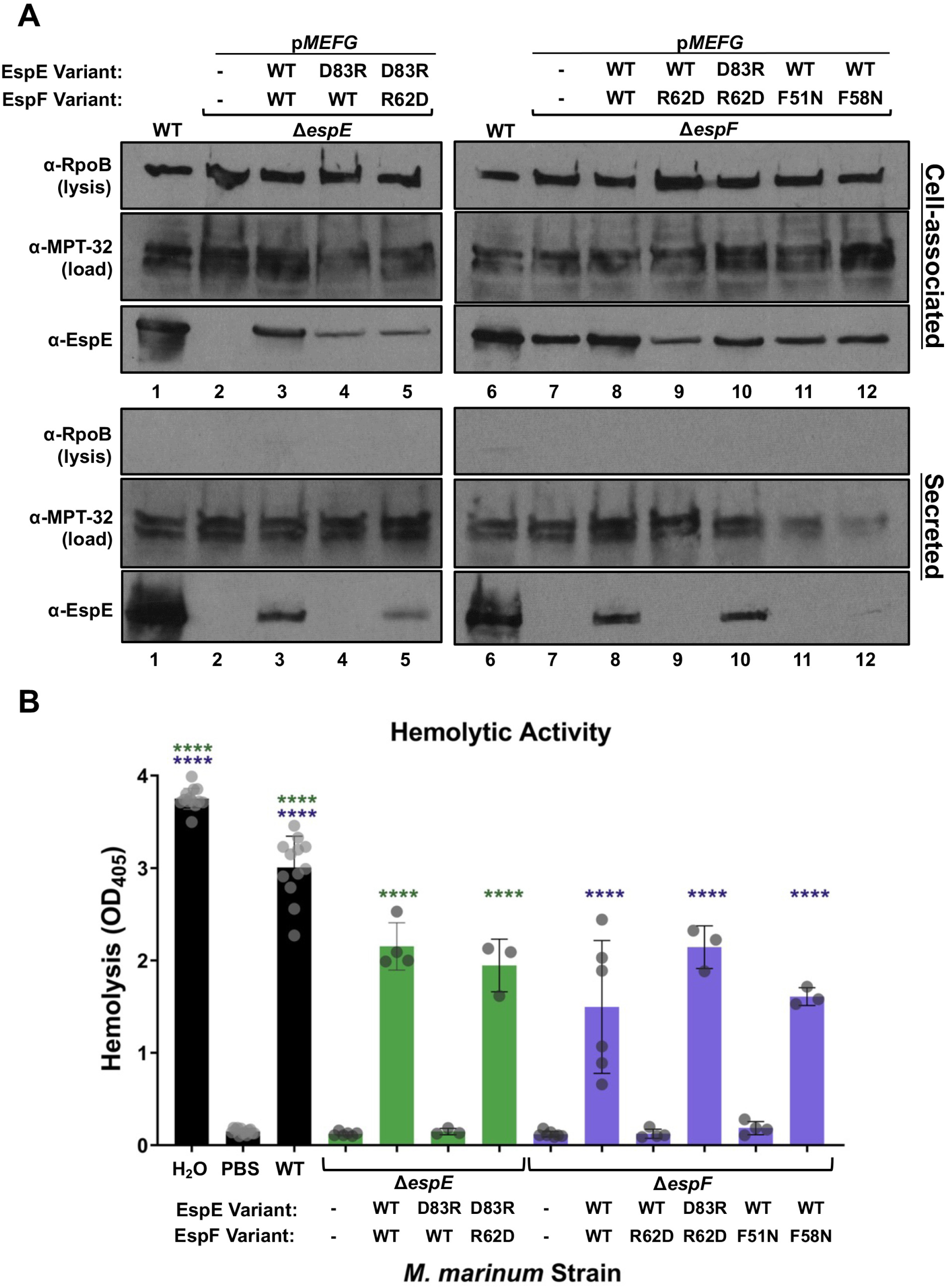
EspE and EspF interaction is required for EspE secretion and hemolytic activity. A) Immunoblot analysis of cell-associated (upper) and secreted (lower) protein fractions from *M. marinum* strains expressing *espE* and *espF* variants in the Δ*espE* (right) and Δ*espF* (left) backgrounds. Data shown are representative of three biological replicates. B) Hemolytic activity of *M. marinum* strains. Data presented are the average of at least three biological replicates in technical triplicate. Statistical significance was determined using a one-way ordinary ANOVA (*P* < 0.0001) followed by Dunnett’s multiple comparison test relative to the Δ*espE* strain (green) or the Δ*espF* strain (purple). **** *P* < 0.0001.

**Table 1.** EspF/ EsxA Interaction Summary.

| Allele | EspE secretion | EsxBA secretion | Hemolysis | Regulation |
| --- | --- | --- | --- | --- |
| EspFL5R | Low | WT | Low | WT |
| EspFV7R | Undetectable | WT | Non-hemolytic | WT |
| EspFD91R | Undetectable | WT | Non-hemolytic | WT |
| EsxAM83I | Undetectable | Undetectable | Non-hemolytic | WT |
| EsxAM93T | Low | low | WT | WT |

We next tested if disruption of the salt bridge affected EspE secretion from the Δ*espF* strain (Fig. 3A, right). EspE was detected in all the Δ*espF* strains (Lanes 6-12, upper). EspE was secreted from the WT strain, but not from the Δ*espF* strain (lanes 6-7, lower), consistent with our prior work (13). Introduction of the WT *espF* gene, but not the *espFR26D* gene, in the Δ*espF* strain restored EspE secretion (lanes 8 and 9). However, as in the Δ*espE* background, the introduction of both the *espED83R* and *espFR62D* genes in the Δ*espF* strain restored EspE secretion (lane 10). Therefore, the reversal of this salt bridge restored secretion of EspE. From these data, we conclude that the EspED83 and EspFR62 residues, which are predicted to participate in a salt bridge, are required for the secretion of EspE and EspF from *M. marinum*.

To further examine the predicted protein-protein interaction between EspE and EspF, we tested if disruption of the large hydrophobic residues of EspF affected EspE secretion (Fig. 1E). Expression of the *espF* variants did not measurably affect EspE levels by immunoblot analysis (Fig. 3, lanes 11 and 12). Introduction of the *espFF51N* gene in the Δ*espF* strain did not promote detectable EspE secretion (lane 11, lower). However, introduction of the *espFF58N* gene the Δ*espF* strain resulted in low, but detectable levels of EspE secretion (lane 12). These data suggest that the hydrophobic residues located further from the salt bridge have increased contribution to the hydrophobic effect, which mediates the predicted interaction between EspE and EspF.

The contact-dependent hemolytic activity of *M. marinum* is dependent on ESX-1 secretion (33) and is therefore often used as a proxy for a strain’s ability to lyse the phagosomal membrane (34, 35). We hypothesized that the *M. marinum* strains expressing EspE or EspF variant proteins which disrupted EspE secretion would be non-hemolytic. As shown in Figure 3B, WT *M. marinum* was hemolytic. Distilled water and phosphate buffered saline (PBS) are cell-free controls for maximum and minimal lysis. Deletion of either the *espE* or *espF* genes resulted in a loss of hemolytic activity, consistent with our previous findings (9, 13, 33). Hemolysis was restored upon expression of the WT *espMEFG* genes, as well as introduction of both the *espE*D83R and *espFR62D* genes in the Δ*espE* or Δ*espF* strains restored hemolysis to similar levels. Likewise, introduction of the *espFF58N* gene in the Δ*espF* strain resulted in hemolytic activity similar to the complemented strain. Introduction of the *espED83R* gene in the Δ*espE* strain, or the *espFR62D* or *espFF51N* genes in the Δ*espF* strain, did not restore hemolytic activity, which was not significantly different from the deletion strains. These data demonstrate that residues predicted to be important for EspE and EspF interaction, which are important for secretion, are also required for hemolytic activity. In addition, the low levels of EspE secretion supported by the introduction of the *espFF58N* gene in the Δ*espF* strain is sufficient to mediate hemolytic activity. Taken together, our data support that EspE and EspF interact through the predicted salt bridge and hydrophobic interface.

### EsxA-EspF interaction is required for secretion and hemolytic activity

The N-termini of EsxA and EspF are predicted to form a two-stranded β-sheet at the interface of the predicted EspE, EspF, EsxA, and EsxB interaction (Fig. 1). As shown in Fig. 1B, the EspF β-strand contains two interior-facing hydrophobic residues, Leu5 and Val7, which are further stabilized by, or acting to stabilize, EspFY87 and EsxAW6. To destabilize this region, we introduced bulky, hydrophilic arginine residues to generate the EspFL5R and EspFV7R protein variants. Across from the EspF-EsxA β-sheet, EspFD91 is predicted to form a network of hydrogen bonds with residues in both EsxA and EsxB. We hypothesized that a charge reversal in this site would sufficiently destabilize this hydrogen bonding network in the EspFD91R variant. Further, EsxAM83 is involved in this interface, and the side chain is positioned directly between the EspFD91 and EsxAW6 side chains. EsxAM93 is solvent-exposed and is slightly outside of this region (Fig. 1C). *M. marinum* strains expressing the *esxAM83I* and *esxAM93T* genes were previously reported to have reduced hemolytic activity and phagosomal damage, but still secreted the EsxA and EsxB substrates (36). We predicted that if the interaction between EspF and EsxA is required for the secretion of the EspE and EspF substrates, then the disruption of the EspF-EsxA interface would prevent EspE secretion, but allow EsxA and EsxB secretion. To test this hypothesis, we generated point mutations in the integrating p*MEFG* and p*esxBA* plasmids. We verified each mutation by targeted Sanger sequencing and introduced the resulting plasmids into *M. marinum* strains containing unmarked deletions of the *espF* gene or *esxBA* locus as appropriate (Fig. 2C and D). We next performed RT-qPCR to verify that the *esxBA* mutant strains produced *esxB* transcript similarly to the complement strain (Fig. 2F). *esxB* transcript was produced in the WT strain, but not the Δ*esxBA* strain. The introduction of the p*esxBA* wild-type and mutant plasmids produced similar levels of *esxB* transcript.

We characterized the functionality of the mutant proteins in promoting ESX-1 secretion by generating cell-associated and secreted protein fractions from *M. marinum* grown in laboratory culture. Our data are summarized in Table 2. We separated the proteins using gel electrophoresis and measured EspE and EsxB levels using immunoblot analysis. As shown in Figure 4, the EspE protein was detected in all Δ*espF* strains (Fig. 4A, lanes 1-6, upper, and Fig. S1A). EspE was secreted from the WT strain, but not from the Δ*espF* strain (lanes 1 and 2, lower). Introduction of the WT *espF* gene in the Δ*espF* strain restored EspE secretion (lane 3, lower). Introduction of the *espFL5R*, *espFV7R, and espFD91R* genes in the Δ*espF* strain resulted in production of the EspE protein. EspE secretion was not detected from strains expressing the *espFV7R* and *espFD91R* genes (lanes 4 and 5, lower). EspE secretion from the Δ*espF* strain expressing the *espFL5R* gene was detectable at very low levels in some samples by immunoblot analysis (Fig. S1A lane 2, arrow). EsxB was produced to reduced levels in the Δ*espF* strain expressing the WT *espF* and *espFL5R* genes (lanes 3 and 4, upper). All strains secreted EsxB (lanes 1-6, lower). These data indicate that the residues predicted to mediate the interaction between EsxA-EspF are required for EspE secretion, but are dispensable for EsxB secretion.

**Figure 4.**
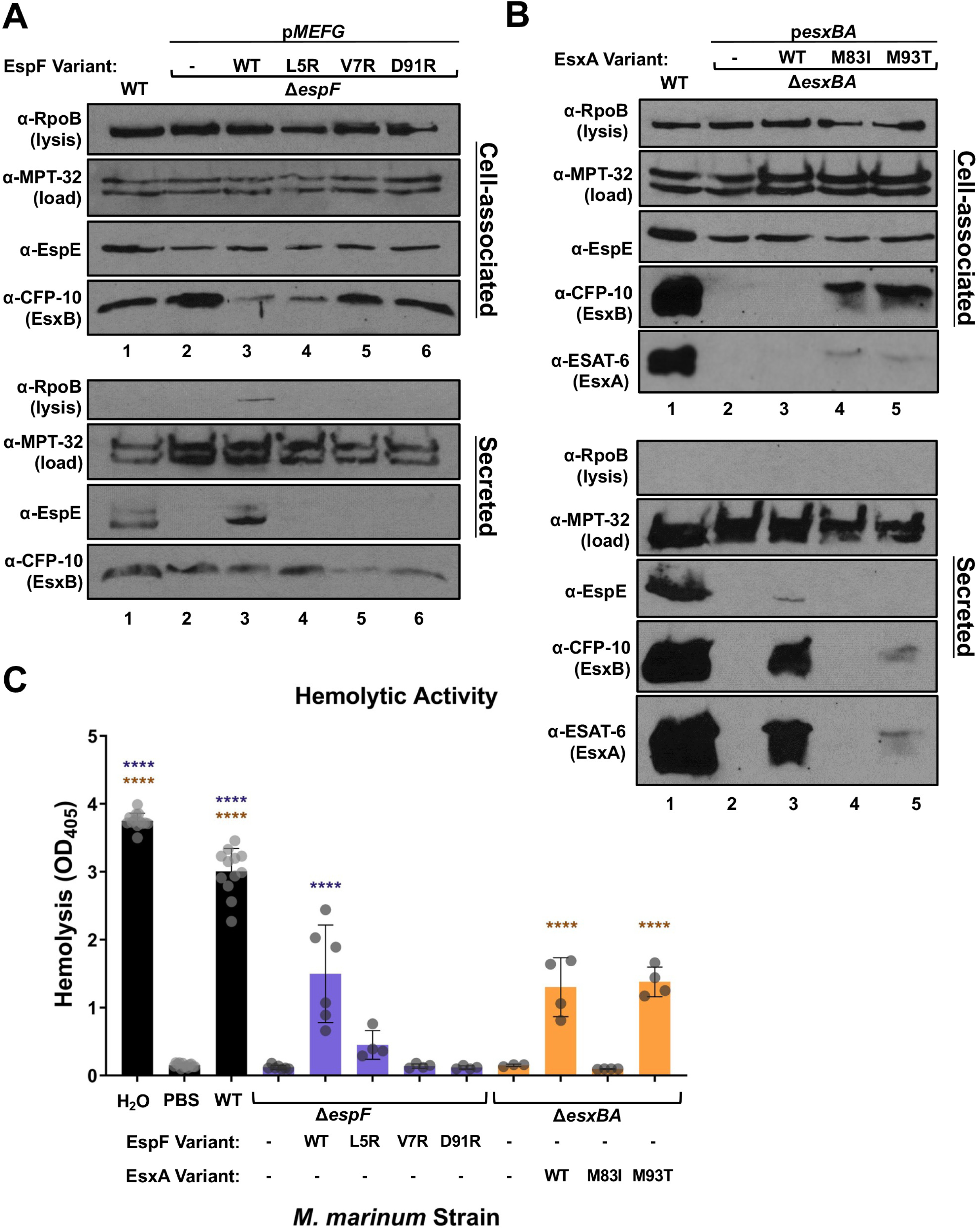
EspF interaction with EsxA is required for EspE secretion. A, B) Immunoblot analysis of cell-associated (upper) and secreted (lower) protein fractions from *M. marinum* strains expressing *espF* (A) and *esxA* (B) mutants. Data shown are representative of three biological replicates. 10μg protein loaded per lane. C) Hemolytic activity of *M. marinum* strains. Data presented are the average of at least three biological replicates in technical triplicate. Statistical significance was determined using a one-way ordinary ANOVA (*P* < 0.0001) followed by Dunnett’s multiple comparison test relative to the Δ*espF* strain (purple) or Δ*esxBA* strain (yellow). **** *P* < 0.0001. Error bars indicate standard deviations.

**Table 2.** EspE/EspF Interaction Summary.

| Allele | EspE secretion | Hemolysis | Regulation |
| --- | --- | --- | --- |
| EspED83R | Undetectable | Non-hemolytic | WT |
| EspFR62D | Undetectable | Non-hemolytic | WT |
| EspED83R<br>EspFR62D | WT | WT | WT |
| EspFF51N | Undetectable | Non-hemolytic | WT |
| EspFF58N | Low | WT | WT |

We next tested if ESX-1 secretion was affected by the introduction of the *esxAM83I* and *esxAM93T* mutant genes. The EsxAM83 and EsxAM93 residues are not predicted to directly participate in the EsxA-EsxB protein-protein interface. Instead, these residues may be involved in the predicted interaction of EsxA with EspF (Fig. 1C). We hypothesized that the reduction in hemolytic activity of strains expressing the *esxAM83I* and *esxAM93T* genes as observed in Osman et al (36) was due to reduced EspE/EspF secretion. EsxB was produced in all but the Δ*esxBA* strain (Fig. 4B, upper; Fig. S1B). Notably, EsxA and EsxB levels and *esxB* transcript levels were significantly decreased in the complementation strain compared to the WT strain (Fig. 4B, 2F). EsxA and EsxB require each other for stability and secretion (14, 15, 37). As such, EsxB and EsxA were secreted from the WT strain but not from the Δ*esxBA* strain (lanes 1-2). Introduction of WT *esxBA* in the Δ*esxBA* strain restored EsxB and EsxA secretion (lane 3, lower). Introduction of the *esxAM83I* gene in the Δ*esxBA* strain was unable to promote EsxB or EsxA secretion (lane 4, and Fig. S1B, lane 4). Introduction of the *esxAM93T* allele in the Δ*esxBA* strain resulted in reduced EsxB and EsxA secretion compared to the complementation strain (lane 5, and Fig. S1B, lane 5). EspE was produced at similar levels in all strains (Fig. 4B, upper). Expression of WT *esxBA* in the Δ*esxBA* strain restored EspE secretion, albeit to levels lower than the WT strain (lane 3), consistent with the reduced secretion of EsxA and EsxB. Secretion of EspE from the Δ*esxBA* strain expressing the *esxAM83I* gene was not detectable (lane 4, and Fig S1B lane 4). The secretion of EspE from the Δ*esxBA* strain expressing the *esxAM93T* gene was lower than the complemented strain, but detectable by immunoblot analysis (lane 5, lanes 4-5; Fig. S1B). These data indicate that the EsxAM83I and EsxAM93T protein variants promoted reduced EsxA and EsxB secretion. The EsxAM93T protein supported low levels of EspE secretion, while the EsxAM83I protein did not support detectable EspE secretion.

Based on the findings above, we predicted that the strains with undetectable EspE secretion would be nonhemolytic. As shown in Figure 4C, the WT *M. marinum* strain was hemolytic. Deletion of the *espF* or *esxBA* genes led to a loss of hemolytic activity, which was partially restored by introduction of the WT genes in the Δ*espF* or the Δ*esxBA* strains (*P* < 0.0001, compared to the deletion strains). The *ΔespF* strain expressing the *espFV7R* or *espFD91R* genes, and the Δ*esxBA* strain expressing the *esxM83I* gene were non-hemolytic, similar to the parental deletion strains, consistent with the loss of detectable EspE secretion. The Δ*espF* expressing the *espFL5R* gene showed reduced hemolytic activity, but the level did not reach significance from the Δ*espF* strain, matching the low, and sometimes undetectable, level of EspE secretion from this strain. Despite the low level of EspE secretion, the Δ*esxBA* strain expressing the *esxAM93T* exhibited hemolytic activity similar to the complementation strain and significantly different from the Δ*esxBA* strain (*P* < 0.0001). These data indicate that the disruption of small hydrophobic residues, or the hydrogen bonding network near the predicted β-strand disrupted ESX-1 function. However, the EsxAM93T variant, which is predicted to disrupt solvent-exposed residues away from the β-strand interaction site, restored ESX-1 function to the Δ*esxBA* strain. Interestingly, this strain did show decreased EsxB secretion compared to the complementation strain, indicating that solvent-exposed EsxAM93 affects EsxA-EsxB protein function.

### The interactions between EspF and EspE or EsxA are dispensable for regulation

In addition to functioning as secreted virulence factors, EspE and EspF negatively control the transcription of the *whiB6* gene (9). WhiB6 is a positive transcriptional regulator of several ESX- 1 substrate genes (19–21). We tested if the residues required for the predicted EspE/EspF and EspF/EsxA interactions are required for regulation of *whiB6* transcription. We performed RT-qPCR to measure *whiB6* transcript levels in these strains. As shown in Fig. 5A, average *whiB6* transcript levels in the Δ*espE* strain were increased, but not significantly different from, compared to the WT strain (*P* = 0.5294). *whiB6* transcript levels were significantly higher in both the WT (*P* = 0.0077) and the Δ*espE* strain (*P* < 0.0001) relative to the Δ*espE* strain expressing the wild-type *espE* gene. The introduction of the *espED83*R gene, or both the *espED83R* and *espFR62D* genes in the Δ*espE* strain resulted in *whiB6* transcript levels that were not significantly different from the expression of the wild-type *espE* gene in the Δ*espE* strain. These data suggest that the EspE variants which impacted secretion and hemolytic activity (Fig. 3) functionally downregulated *whiB6* transcription similarly to the WT EspE protein. In the Δ*espF* strain, *whiB6* transcript levels were significantly increased compared to the WT strain (*P* < 0.0001). However, introduction of the wild-type *espF* gene, the *espF*R62D gene, or the *espE*D83R and *espF*R62D genes together in the Δ*espF* strain, resulted in *whiB6* transcript levels that were not significantly different from expression of the WT strain. Together, these data suggest that the residues predicted to mediate an interaction between EspE-EspF are dispensable for the negative regulation of *whiB6* gene expression.

**Figure 5.**
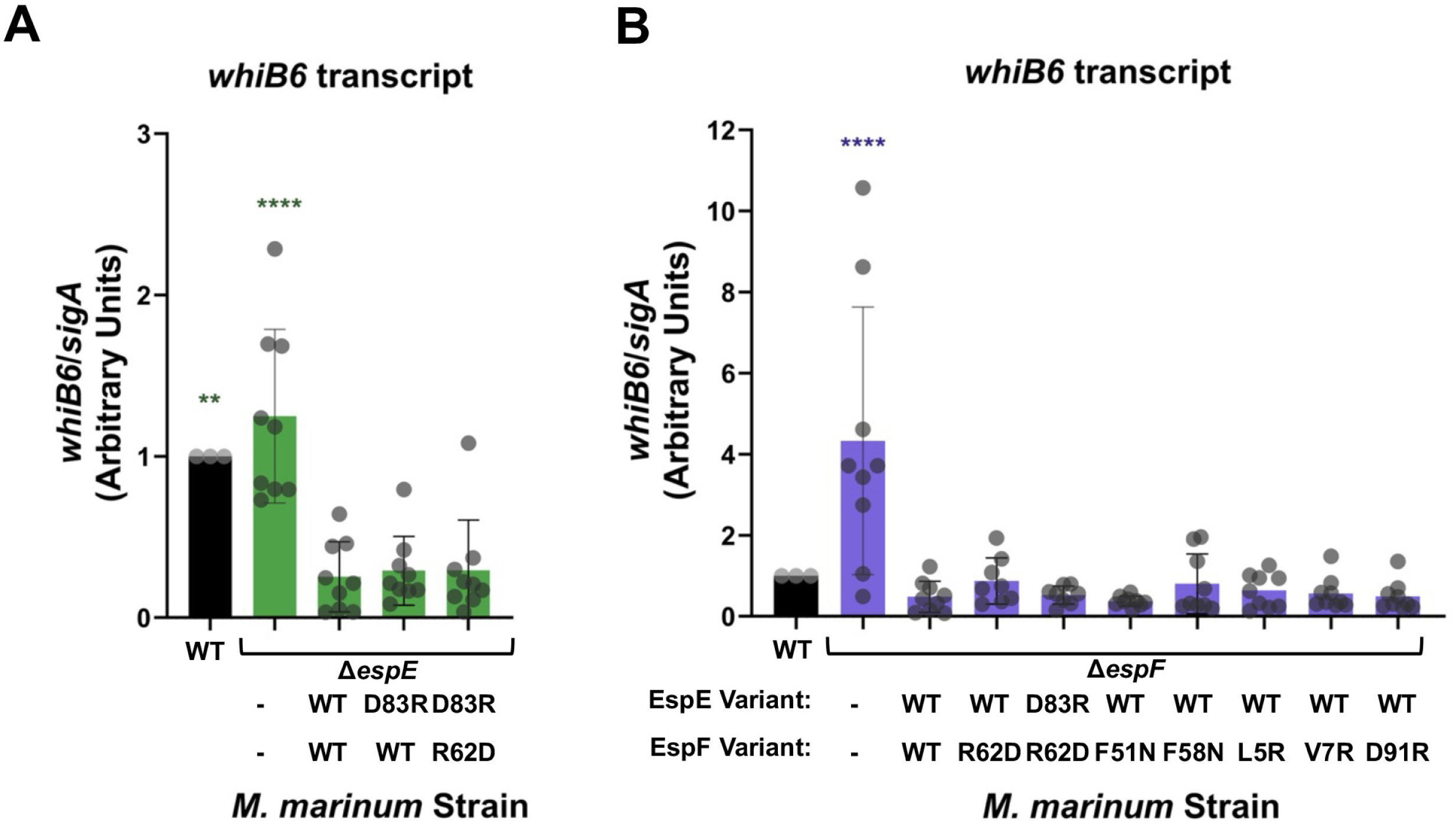
**Neither EspE and EspF interaction nor EspF and EsxA interaction is required for regulation of *whiB6.*** A, B) RT-qPCR analysis of *whiB6* transcript relative to *sigA* transcript in the indicated *M. marinum* strains. ΔΔC_T_ compared to the WT strain is shown. Data shown are three biological replicates each with three technical replicates. Statistical significance was determined using a one-way ordinary ANOVA (*P* < 0.0001) followed by Dunnett’s multiple comparison test relative to the Δ*espE/*p*MEFG* strain (A) or Δ*espF/*p*MEFG* strain (B). ** *P* = 0.0077, **** *P* < 0.0001. Error bars indicate standard deviations.

We next tested if the residues predicted to mediate protein-protein interaction between EspF and EsxA were essential for the negative regulation of *whiB6* gene expression by EspF. We performed RT-qPCR to determine relative levels of *whiB6* transcript in these strains. As shown in Figure 5B, introduction of the *espFL5R*, *espFV7R*, and *espFD91R* alleles in the Δ*espF* strain resulted in *whiB6* transcript levels comparable to, and not significantly different from, the Δ*espF* strain expressing the wild-type *espF* gene. All strains expressing *espF* resulted in significant repression of *whiB6* transcript levels relative to the Δ*espF* strain (Dunnett’s Multiple Comparisons test, *P* < 0.0001). From these data, we conclude that the residues predicted to promote the interaction between EspF and EsxA are dispensable for the regulatory function of EspF.

WhiB6 is a positive regulator of ESX-1. Upon increased expression of *whiB6,* there is an associated increase in ESX-1 substrates, including EsxB (19, 20). We hypothesized that the levels of *whiB6* transcript in the Δ*espF* strains expressing the *espF* variants would reflect the levels of EsxB protein as detected by immunoblot. As shown in Fig. 4A, EsxB protein was increased in the Δ*espF* strain compared to the WT strain and the complementation strain, consistent with our previously published data (9). In all strains expressing mutant *espF* alleles, EsxB protein was decreased relative to the Δ*espF* strain. These data support that the residues predicted to promote the interaction of EspF with EspE or EsxA are dispensable for the negative regulation of *whiB6* transcription by EspE and EspF.

## Discussion

In this study, we used structural modeling to predict residues important for the interaction between the ESX-1 substrates EspE/EspF, and the interaction of this protein pair with the EsxA/EsxB heterodimer. Our computational model served as a foundation to make targeted mutations to probe the importance of these protein-protein interfaces. We tested the requirement of these residues in protein secretion, hemolytic activity, and transcriptional regulation. Our results show that the residues likely involved in interaction between EspE and EspF, and for EspF with EsxA are important for protein secretion and for hemolytic activity, but are dispensable for regulatory function of these substrates. EspE and EspF require each other for secretion, and are secreted after the other ESX-1 substrates (13). However, to our knowledge, no previous studies have shown evidence that EspE and EspF directly interact with one another. In addition, the mechanisms of ESX-1 substrate transport through the mycobacterial cell wall are largely unknown. We provide support for a model in which EspE and EspF interact, and that this interaction is required for their secretion (Fig. 6, “A”). Further, we suggest that EspF and EsxA interaction facilitates either targeting (Fig 6, “A”) transport (Fig. 6, “B”) of the EspE/EspF substrate pair to the extracellular environment. If the interactions between EspF and EsxA promote transport across the cell envelope, it is likely that EspF or EspE also interact with other substrates, including the EspB, EspJ or EspK group II substrates (Fig. 6, “C”). Finally, we show that the protein-protein interactions of EspE and EspF are dispensable for regulation of *whiB6* transcription in the cytoplasm (Fig. 6, “D”).

**Figure 6.**
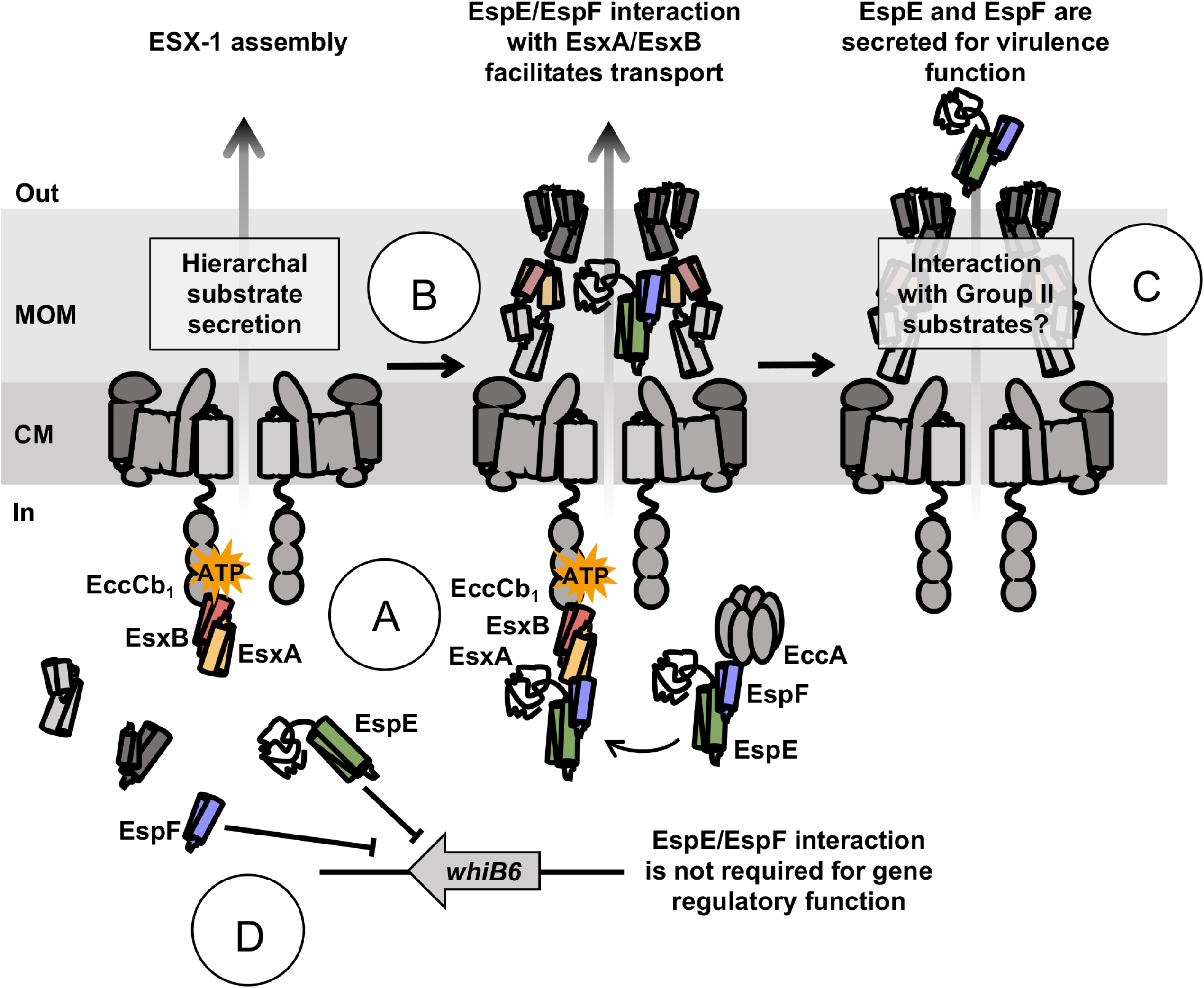
Model of EspE/EspF protein-protein interaction in relation to secretion and gene regulation. Our work supports a model in which EspE/EspF interaction is separable from their regulatory functions. The EsxA/EsxB heterodimer interacts with the EccCb_1_ ATPase, thereby targeting the ESX-1 substrates for secretion (A). EspF interacts with EccA (23), and this interaction may function to transport the EspE/EspF heterodimer to the EsxB/A dimer and the ESX-1 membrane complex (A). The interaction of EspE/EspF with EsxA/EsxB facilitates either targeting or transport through the mycobacterial cell wall via an unknown mechanism (B). Based on our prior model of hierarchical secretion (13), the EspE/EspF heterodimer may interact with other ESX-1 substrates during transport (C). Within the cell, EspE and EspF perform their regulatory functions separately (D).

A common theme in ESX-1 protein secretion is that substrates are secreted in pairs. EsxA and EsxB were the first pair of substrates identified (28, 37). In *M. tuberculosis,* EspA and EspC directly interact with one another in the mycobacterial cytosol prior to export (22). We previously showed that deletion of the *espF* gene prevents the secretion of EspE, and the deletion of the *espE* gene prevents the secretion of EspF (13). In *M. tuberculosis* and *M. marinum,* the *espACD* and *espEFG_1_* loci are paralogous (22). Although the residues mediating the EspA/EspC interaction are not known, the paralogous relationship between EspA/EspC and EspE/EspF supports our hypothesis that EspE and EspF interact. Likewise, modeling using AlphaFold2 supports that EspA/EspC also form a PE-PPE-like heterodimer (K. Korotkov, unpublished data). We identified two key interaction points between EspE and EspF: an electrostatic interaction between residues EspED83 and EspFR62, and a hydrophobic, intra- helical-bundle pocket. Our data suggests that the hydrophobic residues positioned further from the electrostatic interaction are essential for EspE-EspF interaction through the intra-helical-bundle pocket. However, electrostatic interactions are more likely to confer specificity in protein interactions (38). Likewise, we provide evidence that the predicted salt bridge between EspED83 and EspFR62 is specific to EspE-EspF interaction. Our data further support that the EspE and EspF are a pair of ESX-1 substrates that directly interact.

ESX-1 substrates are secreted in an ordered hierarchy. In *M. marinum*, PPE68 and MMAR_2894 are secreted prior to the EsxA/EsxB substrate pair (13, 35, 39). EsxB, but not EsxA, interacts with EccCb_1_ through its C-terminus (17, 30, 37), targeting the substrate pair for secretion. EsxA and EsxB are required for the secretion of all other known substrates including EspB, EspK, EspJ, EspE and EspF (12, 13), but the mechanism underlying this requirement is unknown. There is significant support in the literature for a model in which EsxA and EsxB facilitate the transport of ESX-1 substrates from the mycobacterial cytosol by interaction with the EccCb_1_ ATPase. We previously showed that EspF directly interacts with EsxA, EspC, and the EccA_1_ ATPase (23). The direct interaction between EspF and EsxA by both co-immunoprecipitation and the yeast-two-hybrid system supports our hypothesis that the secretion of the EspE/EspF protein pair is mediated by EsxA/EsxB. In this study, we identified that the N-termini of EsxA and EspF are predicted to form a small, two-stranded β-sheet made up primarily of small hydrophobic residues. Adjacent to this region, there is a large hydrogen bonding network that spans the EsxA-EsxB-EspF interface. Mutation to either of these regions led to a decrease or loss of EspE secretion and hemolytic activity, and mutation to this region of EsxA specifically resulted in decreased secretion of EsxA and EsxB. Together, our results support a model whereby protein interaction is required for the targeting and secretion of ESX-1 substrates, while the regulatory activity of EspE and EspF do not require substrate interactions, allowing them to function independently.

Previous studies observed that the EsxAM93T variant promoted reduced EsxA and EsxB secretion from *M. marinum*, and suggested that low levels of EsxA secretion was sufficient for ESX-1-mediated lytic activity (34, 36). The EsxAM83I and EsxAM93T protein variants were first studied by Osman *et al.* (36), and it was suggested that both resulted in a loss of virulence, but retained ESAT-6 (EsxA) and CFP-10 (EsxB) secretion. Aligned with this study, we found that very little secreted EspE protein is necessary for hemolysis. The hemolytic activity of our *M. marinum* strains appears to correlate with the level of secreted EspE protein, rather than that of EsxA (See Table S1 and S2). Indeed, deletion of *espE* does not significantly impact the secretion of EsxA, while EsxA is essential for EspE secretion (12, 13). However, our data shows that introduction of the *esxAM83I* gene in the Δ*esxBA* strain resulted in a loss of EsxB and EsxA secretion and hemolytic activity, whereas introduction of the *esxAM93T* gene in the Δ*esxBA* strain resulted in decreased EsxB secretion but did not impact hemolytic activity. However, our study and the Osman *et al.* study are not necessarily in opposition. Osman *et al.* showed that the expression of *esxAM93T* results in both higher EsxB secretion and average percent hemolysis than the expression of *esxAM83I*, but this level of hemolytic activity did not reach significance compared to the *esxA* deletion strain (36).

We previously demonstrated that EspE and EspF are dual-functioning ESX-1 substrates: both proteins also negatively regulate ESX-1 gene expression in the mycobacterial cytoplasm (9). Deletion of either the *espE* or *espF* gene resulted in increased *whiB6* transcript in *M. marinum* (9). Our current study showed that mutations predicted to disrupt the interaction between EspE and EspF did not impact the regulatory activity or either protein. We expect that our complementation strains exhibited lower levels of *whiB6* transcript compared to the WT strain due to the presence of the additional copy of *espM* and *espF,* both negative regulators of ESX-1 (9, 40), in the *pMEFG* plasmid. In our prior work, we showed that cross-complementation of these genes did not restore of *whiB6* expression to WT levels. In addition, the overexpression of *espE* and *espF* in the WT strain led to differing phenotypes. Together, our data support that EspE and EspF function to regulate *whiB6* independently from one another, which is consistent with EspE and EspF functioning as regulators independent of their interactions. One possibility is that the individual regulatory functions of EspE and EspF are modulated by the direct interaction of these substrates and subsequent secretion from *Mycobacterium*.

We propose a model in which EspE and EspF interact, and EspF-EsxA interaction facilitates the transport of the EspE/EspF substrate pair through the mycobacterial cell wall (Fig. 6). Our research leads to further questions about how EspE and EspF are secreted. It is unknown if the EspE/EspF substrate pair is handed off to other group I or group II ESX-1 substrates during translocation. In addition, it is unknown if EspE-EspF interaction plays a role in virulence following their secretion. Both EsxA and EspF are known to be N-terminally acetylated (41, 42). This N-terminal acetylation is not accounted for in our model of protein-protein interaction. However, N-terminal acetylation of EsxA is dispensable for ESX-1 function in *M. marinum* (43). It is unknown how the N-terminal acetylation of EspF plays a role in substrate transport, virulence, and gene regulation. In addition, while we were able to create a compensatory mutation to support EspE/EspF interaction, we were unable to do the same for EsxA/EspF interaction. It is possible that this set of mutations disrupted EspE/EspF secretion other than through protein-protein interaction with the EsxA/B dimer. These questions encompass future studies to further elucidate the mechanisms of EspE and EspF function. Further studies, including biochemical approaches to demonstrate direct interaction between EspE and EspF, are forthcoming.

Our model does not incorporate the C-terminal half of EspE. The largely disordered C-terminal half of EspE is predicted to contain several alpha-helical regions, which may overlay the EspF binding site (Figure S7). We modeled the full-length EspE protein with two structure prediction pipelines: AlphaFold2 (24, 31) and Robetta (RoseTTAFold) (44). While AlphaFold2 did not predict any ordered interaction between the N-terminal and C-terminal domains, the RoseTTAFold prediction shows EspE’s C-terminal α-helices overlaying the predicted EspF binding site. AlphaFold2 is known to overestimate disorder, which may contribute to the differences between the two models (45). This region may act to stabilize EspE prior to EspF interaction. Figure S4 shows the hydrophobicity of the EspF and EspE C-terminal α-helices at the predicted protein-protein interface. The binding face of EspF extends further down, and contains more hydrophobic residues, than the EspE C-terminal helices. Interestingly, the general pattern of hydrophobic and hydrophilic residues is maintained between the two structures. The relationship between the EspE C-terminal helical structure and protein stability and secretion could be the focus of further study.

In conclusion, we have shown evidence that the dual-functioning virulence factors EspE and EspF have separable functions to regulate the ESX-1 system. We also provide evidence that EspE and EspF interact. This interaction is required for EspE secretion and virulence. However, EspE-EspF interaction is dispensable for the negative regulation of *whiB6,* a positive regulator of ESX-1. This suggests that EspE and EspF function independently in their roles in gene regulation. In addition, EspF and EsxA directly interact, and this interaction is required for EspE secretion and hemolytic activity. We propose that EspF interaction with EsxA facilitates the translocation of EspE and EspF through the mycobacterial cell wall. We were able to use a computational model of ESX-1 substrate interaction to further define the mechanisms of substrate function and transport.

## Materials and Methods

### Generation of DNA constructs

All DNA constructs were generated using oligonucleotide primers purchased from Integrated DNA Technologies (IDT, Coralville, IA). The p*espMEFG* and p_MOP_*esxBA* plasmids were generated as described previously (9, 19),. All constructs were verified using targeted Sanger DNA sequencing performed by the Genomics and Bioinformatics Core Facility at the University of Notre Dame. All strains, plasmids, and oligonucleotide primers are listed in Tables S1, S2 and S3.

### Generation of Point Mutations using Site-Directed Mutagenesis (SDM)

To generate individual point mutations in the *espE, espF,* or *esxA* genes in the p*espMEFG* and p_MOP_*esxBA* plasmids, we designed mutagenic oligonucleotide primers (IDT) according to the guidelines in the QuikChange Site-Directed Mutagenesis Kit Instruction Manual (Agilent). Mutagenesis PCR reactions were performed as described previously(46) using PfuTurbo DNA Polymerase Alternative Detergent (Agilent).

### Growth of bacterial strains

*E. coli* were grown in LB (Luria Bertani) broth (VWR) or on LB agar. Plasmids were maintained in DH5α cells (NEB). *E. coli* strains were grown at 37°C and were supplemented with hygromycin B (Corning or RPI) as necessary (200μg/mL).

All *M. marinum* strains in this study are isogenic and generated from the M parental strain (ATCC BAA-535). *M. marinum* was grown on Middlebrook 7H11 agar base (Millipore Sigma) supplemented with 0.5% glycerol and 0.5% glucose, or in liquid media Middlebrook 7H9 broth base (Millipore Sigma), supplemented with 0.5% glycerol and 0.1% Tween-80 (VWR). *M. marinum* was grown at 30°C and supplemented with hygromycin B (Corning or RPI) as necessary (50μg/mL).

### Hemolysis Assays

Sheep red blood cell (sRBC) hemolysis assays were performed exactly as described previously (19).

### Secretion Assays and Immunoblot Analysis

ESX-1 secretion assays were performed as described previously (47). Briefly, *M. marinum* strains were grown in Sauton’s defined media for 48 hours. Cells were harvested by centrifugation, resuspended in PBS, and mechanically lysed to generate the cell-associated fraction. Secreted fractions were filtered through Nalgene rapid flow 0.2μm filters (Thermo) and concentrated in a centrifugal filter unit. Protein concentrations were determined using the Micro BCA™ Protein Assay Kit (Thermo).

Immunoblotting was performed as previously described (19). Briefly, 10-20μg of protein samples were separated on 12% Acrylamide SDS-PAGE gels or 4–20% Mini-PROTEAN® TGX™ Precast Protein Gels (Bio-Rad). Protein was transferred to nitrocellulose membrane (Cytiva Amersham^TM^ Protran^TM^ 0.2μm NC) at 100V for one hour. Membranes were incubated in 5% milk in PBS Tween-20 with primary antibodies against: Anti-*E. coli* RNA Polymerase β [8RB13] (RpoB, VWR, 1:20,000 dilution), Anti-MMAR_5439 (EspE, Genscript, 1:5000 dilution, (48)). The following reagents were obtained through BEI Resources, NIAID, NIH: Polyclonal Anti-*Mycobacterium tuberculosis* CFP10 (Gene *Rv3874*) (antiserum, Rabbit), NR-13801 (used at a 1:5000 dilution), and Polyclonal Anti-*Mycobacterium tuberculosis* Mpt32 (Gene *Rv1860*) (antiserum, Rabbit), NR-13807 (used at a 1:20000 dilution). Secondary antibodies were used at 1:5000 dilution. Anti-mouse IgG, HRP-linked Antibody (Cell Signaling Technology) was utilized for detection of RpoB. HRP-conjugated goat anti-rabbit IgG secondary antibody (BioRad) was used for detection of EsxB, EspE, and MPT-32. Uncropped immunoblots are in Figures S2-S6, and are representative of at least three independent biological replicates.

### RT-qPCR

Reverse transcriptase quantitative PCR (RT-qPCR) was performed as previously described (40). Briefly, 200ng of RNA was DNase treated (Promega) with the addition of 1μL of 100nM CaCl_2_ and 50nM MgCl_2_. DNase treated RNA was reverse transcribed into cDNA using Superscript II reverse transcriptase according to the manufacturer’s instructions (Invitrogen). cDNA was quantified by Nanodrop (Thermo). RT-qPCR was performed using 250ng of the synthesized cDNA and SYBR Green PCR master mix (Applied Biosystems, Thermo). Oligonucleotide primers are listed in Table S5. A QuantStudio 3 PCR system (Applied Biosystems, Thermo) was used to run each reaction. Data was analyzed to determine the ΔΔC_T_ against the WT strain using the following equation:

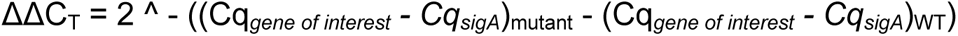

### Statistical Analysis

All statistical analysis was performed using GraphPad Prism 10. In each case the significance was determined using a One-way ANOVA test followed by a Dunnett’s multiple comparison’s test. The *P* value for each experiment is specified in the figure legends and/or text.

### Structural Modeling

Structural models of EspE/EspF in complex with EsxA/EsxB were generated using AlphaFold2-multimer (49) with model weights v.2.2 as implemented in ColabFold (50).

## Acknowledgements

We thank the Champion Lab for their critical reading of this article. The content of this article is solely our responsibility and does not necessarily represent the official views of the National Institutes of Health (NIH). P.A.C. is supported by the National Institutes of Health under award numbers AI156229, AI106872, AI149147, and AI149235. K.V.K. is supported by AI169231. R.J.P., K.V.K. and P.A.C. designed the research; R.J.P. performed the research; R.J.P, K.V.K and P.A.C. analyzed the data and wrote the paper.

